# Amyloid-β exposed astrocytes induce iron transport from endothelial cells at the blood-brain barrier by altering the ratio of apo- and holo-transferrin

**DOI:** 10.1101/2023.05.15.540795

**Authors:** Stephanie L. Baringer, Avraham S. Lukacher, Kondaiah Palsa, Hyosung Kim, Ethan S. Lippmann, Vladimir S. Spiegelman, Ian A. Simpson, James R. Connor

**Author notes:** Correspondence: James R. Connor, Penn State College of Medicine, 500 University Drive, Hershey, PA 17033. **Author Contributions**: SLB, IAS, and JRC designed research. SLB, ASL, and KP performed research, HK, ESL, and VSS contributed reagents. SLB and JRC analyzed data. SLB wrote the paper. **Conflict of interest statement**: The authors declare no competing financial interests.

## Abstract

Excessive brain iron accumulation is observed in early in the onset of Alzheimer’s disease, notably prior to widespread proteinopathy. These findings suggest that increases in brain iron levels are due to a dysregulation of the iron transport mechanism at the blood-brain barrier. Astrocytes release signals (apo- and holo-transferrin) that communicate brain iron needs to endothelial cells in order to modulate iron transport. Here we use iPSC-derived astrocytes and endothelial cells to investigate how early-disease levels of amyloid-β disrupt iron transport signals secreted by astrocytes to stimulate iron transport from endothelial cells. We demonstrate that conditioned media from astrocytes treated with amyloid-β stimulates iron transport from endothelial cells and induces changes in iron transport pathway protein levels. The mechanism underlying this response begins with increased iron uptake and mitochondrial activity by the astrocytes which in turn increases levels of apo-transferrin in the amyloid-β conditioned astrocyte media leading to increased iron transport from endothelial cells. These novel findings offer a potential explanation for the initiation of excessive iron accumulation in early stages of Alzheimer’s disease. What’s more, these data provide the first example of how the mechanism of iron transport regulation by apo- and holo-transferrin becomes misappropriated in disease to detrimental ends. The clinical benefit from understanding early dysregulation in brain iron transport in AD cannot be understated. If therapeutics can target this early process, they could possibly prevent the detrimental cascade that occurs with excessive iron accumulation.

**Significance Statement:** Excessive brain iron accumulation is hallmark pathology of Alzheimer’s disease that occurs early in the disease staging and before widespread proteinopathy deposition. This overabundance of brain iron has been implicated to contribute to disease progression, thus understandingthe mechanism of early iron accumulation has significant therapeutic potential to slow to halt disease progression. Here, we show that in response to low levels of amyloid-β exposure, astrocytes increase their mitochondrial activity and iron uptake, resulting in iron deficient conditions. Elevated levels of apo (iron free)-transferrin stimulate iron release from endothelial cells. These data are the first to propose a mechanism for the initiation of iron accumulation and the misappropriation of iron transport signaling leading to dysfunctional brain iron homeostasis and resultant disease pathology.

## Introduction

Alzheimer’s disease (AD) is a progressive neurodegenerative disease that is characterized clinically by memory impairment and cognitive decline and pathologically by amyloid-β (Aβ) plaques and neurofibrillary tau tangles. In addition to these hallmark pathologies, excessive brain iron accumulation is repeatedly observed in AD patients^1–4^. Unique patterns of regional brain iron accumulation^1, 5^ correlate with disease progression and can reliably predict cognitive impairment in AD patients^2^. Recently, Ayton *et al.* have shown that brain iron accumulation occurs early in AD and prior to widespread Aβ or tau pathology distribution^1^, suggesting iron uptake dysfunction occurs independent from the vascular damage Aβ can inflict in later stages of the disease^6^. Anemic and iron transport processes are also upregulated in AD patient brain tissue^7^, proposing the hypothesis that the AD brain may operate in functional iron deficiency.

Brain iron uptake is tightly regulated by endothelial cells (ECs) of the blood-brain barrier (BBB)^8–11^. Our group and others have shown that astrocytes release signals that modulate iron release from ECs^9, 12^. Importantly, the iron status of astrocytes modulates iron release from ECs based on astrocytic iron needs: iron depleted astrocytes stimulate iron release whereas iron saturated astrocytes suppress iron release^9^. A mechanism for regulating iron release is the ratio of apo-(iron free) to holo-(iron bound) transferrin (Tf) in the extracellular fluid to increase and decrease iron release, respectively^8, 11, 13^. We have also demonstrated this mechanism *in vivo* in healthy mice^11^. In this study we address if the model of apo- and holo-Tf ratio regulating iron release can be misappropriated in a disease setting. Numerous neurological diseases display alterations in brain iron levels that lead or contribute to pathology and symptoms^14^, but it has remained unknown if altered iron levels were due to a dysregulation of iron transport mechanisms.

The majority of iron and AD research has focused on how iron can accelerate and contribute to Aβ and tau pathology deposition^15^. Despite the clear correlation between AD progression and brain iron accumulation, there has been a lack of attention in deciphering causes for increased iron uptake into the brain. In the present study, we offer a model for iron uptake initiation by the way of dysregulation in iron release signals. We demonstrate that astrocytes treated with low levels of Aβ increase their iron uptake, resulting in an increased release of apo-Tf which is a signal of iron deficiency and thus stimulates iron release from ECs. These results provide significant insight into mechanism of iron uptake regulation in response to early AD pathology.

## Materials and Methods

### Cell Culture

Human brain endothelial-like cells (ECs) were differentiated from ATCC-DYS0100 human iPSCs as described previously^16, 17^. Briefly, iPSCs were seeded onto a Matrigel-coated plate in E8 medium (Thermo Fisher Scientific, 05990) containing 10 µM ROCK inhibitor (Y-27632, R&D Systems, 1254) at a density of 15,000 cells/cm^2^. The iPSC differentiation was initiated by changing the E8 medium to E6 medium (Thermo Fisher Scientific, A1516401) after 24 hours seeding. E6 medium was changed every 24 hours and cells were maintained in E6 medium up to 4 days. After 4 days, cells were switched to human endothelial serum free medium (hESFM) (Thermo Fisher Scientific, 11111) supplemented with 10nM bFGF (Fibroblast growth factor, Peprotech, 100-18B) and 10 µM all-trans retinoic acid (RA, Sigma, R2625) and 1% B27 (Thermo Fisher Scientific, 17504-044). Medium was not changed for 48 hours. After 48 hours, cells were collected and replated onto Transwell filters coated with collagen IV and fibronectin. Twenty-four hours after replating, bFGF and RA were removed from the medium to induce barrier phenotype.

iPSC-derived astrocytes were generated using the CC3 iPSC line as described perviously^18^. The undifferentiated iPSCs were cultured in E8 medium on 6-well plates coated with growth factor-reduced Matrigel (Corning). When the iPSCs reached 60-80% confluency, they were subcultured using Versene (Thermo Fisher Scientific). For differentiation, the iPSCs were dissociated using Versene and seeded in low-attachment plates to form embryoid bodies (EBs) with a density of 4×10^5^ cells/well in E6 medium supplemented with 100x N2 and 10 μM Y27632. Neural differentiation in the EBs was achieved through dual inhibition of SMAD signaling for 7 days. Subsequently, the EBs were seeded onto Matrigel-coated plates and cultured for another 7 days. During this time, NPCs were manually isolated and expanded as neurospheres in suspension culture for 7 days. The neurospheres were then dissociated into single cells and seeded onto Matrigel-coated plates for directed astroglial differentiation for 30 days. Medium changes were done every 48 hours with astroglia medium containing 10 ng/mL BMP-4 (PeproTech, 120-05ET), 10% N2 supplement (ThermoFisher, 17502-048), 20% B27 supplement (ThermoFisher, 12587-010), 20 ng/mL bFGF (PeproTech, 100-18B), and 10% penicillin-streptomycin (Hyclone, SV30010), and the cells were passaged at approximately 80% confluency.

### Aβ Peptide Preparation and Treatment

Aβ_42_ was prepared as previously described^19, 20^. Briefly, chilled hexafluoroisopropanol (HFIP) was added to Aβ_42_ peptide (Alfa Aesar, J66387) to obtain a concentration of 1 mM. The peptide solution was sonicated for 5 minutes and incubated at room temperature for 30 minutes. The solution was divided into single use aliquots. HFIP was removed via overnight evaporation and further exsiccated in vacuo in a lyophilizer. The tubes were stored at - 80 °C. To use, the peptide film was reconstituted in corresponding cell culture media with sonication for 10 minutes. Cells were treated with 50 nM Aβ_42_ for all experiments.

### Radiolabeling

^55^Fe (Perkin Elmer) was complexed with 1 mM nitrilotriacetic acid (NTA), 6 mM ferric chloride (FeCl_3_), and 0.5 M sodium bicarbonate (NaHCO_3_) at a ratio of 100 μL NTA: 6.7μL FeCl_3_: 23.3 μL NaHCO_3_: 50 μCi ^55^FeCl_3_ to form the ^55^Fe-NTA complex^11^. After complexing, ^55^Fe-NTA was either used immediately or incubated with apo-Tf (Sigma) for 30 minutes to allow for iron loading. Unbound iron was separated from the total complex using PD midiTrap-G25 columns following manufacturer’s instructions (GE Healthcare Bio-Sciences).

### ^55^Fe-Tf Transport Studies

The radiolabeled iron transport studies were described previously^17^. Initially, the apical chamber of 12-well Transwell plates (Costar Transwell, 0.4 μm pore, Corning) was coated with collagen IV (Sigma) and fibronectin (Sigma) at a ratio of 5:4:1 of ddH2O, 1 mg/ml collagen IV, and 1 mg/ml fibronectin respectively. A total of 200 μl was used to coat plates 4 h at 37 °C. Following coating, the differentiated ECs were replated onto the coated filters. The basal chamber was filled with 1.5 mL of the same media. After allowing the cells to attach overnight at 37 °C, the media was changed in both chambers to hESFM, supplemented with 1% B27, but lacking bFGF and RA. Cells were incubated at 37 °C overnight to allow growth and tight junction formation, after which all transport studies were performed. Before experimental media addition, trans endothelial electrical resistance (TEER) measurements were taken using an Epithelial Volt/Ohm Meter (EVOM2, STX2, World Precision Instruments). Blank (media only) TEER readings were obtained and subtracted from all other TEER measurements. Across all experimental conditions, we report an average TEER value of 3800 ± 126 Ω × cm^2^. At the beginning of the experiment, all media was removed. 500 μl of serum-free media were added to the apical chamber containing,10 μCi of ^55^Fe-Tf (1 mg/ml) and 1 mg/ml RITC-Dextran (70 kD, Sigma) – to monitor tight junction formation and barrier. In the basal chamber, 1.8 mL of the experimental media was placed. At hours 0, 4, 8, and 24, 100 μl was removed from the basal chamber and added to scintillation vials along with 10 mL CytoScint scintillation cocktail (MP Biosciences). Samples were counted using the Hidex 300 SL (LabLogic) for three minutes each. Blank tube values were subtracted from final counts to correct for background counts. At hours 0, 4, 8, and 24, an additional 100 μl was removed from the basal chamber to measure RITC fluorescence (excitation: 555 nm, emission: 580 nm) on a SpectraMax Gemini EM plate reader (Molecular Devices).

### Protein Detection in Media

Frozen aliquoted media samples were thawed for protein detection. Aβ_42_ levels were measured by enzyme-linked immunosorbent assay (Invitrogen, KHB3441). Soluble amyloid precursor protein α (sAPP-α) levels were measured by enzyme-linked immunosorbent assay (IBL, 27734). Interleukin-6 (IL-6) levels were measured by enzyme-linked immunosorbent assay (R&D Systems, D6050). Lactate dehydrogenase (LDH) levels were measured using Cytotoxicity Detection Kit (Roche, 11644793001) according to manufacturer’s directions.

The ratio of apo-to holo-Tf was determined using a urea gel shift assay similar to previously described methods^21^. Briefly, urea binds to the unoccupied iron binding sites on apo-Tf, which results in apo-Tf running slower through a gel and separating from holo-Tf. Conditioned media was concentrated 10x using Amicon Ultra-15 centrifugal filter units 10kD (Millipore, UFC901024). To pull down Tf from the media, 10 μl of anti-transferrin antibody (Abcam, ab82411) was incubated with 100 μl Dynabeads Protein A (Thermo, 10001D). After washing, 100 μl of concentrated media sample was added and allowed to incubate with rotation overnight. The next day, after washing, Tf was eluted from the magnetic beads using 200 μl urea-based elution buffer (8 M urea, 20 mM Tris, pH 7.5, and 100 mM NaCl) with gentle agitation three times. The eluted solution was then concentrated 20x using Amicon Ultra-0.5 centrifugal filer units 50 kD (Millipore, UFC505024). The resulting concentrate was mixed in equal part with TBE urea sample buffer (Bioworld 10530025-1) and loaded on a 10% 15-well TBE urea gel (Biorad 4566036). The gel was run at 170V for 5 hours, after which the gels was washed with water and all protein was stained using RAPIDstain (Calbiochem, 553215) for 1 hour. After additional washing, the stained gel was imaged on an Amersham Imager 600 (GE Amersham). To quantify the percentage of apo-Tf present in samples, a standard curve was made with percentages of apo- and holo-Tf totaling 5 μg of protein. The band intensities of apo- and holo-Tf were determined and the ratio of apo:holo was calculated. The ratio value was plotted against the percentage of apo-Tf to create the standard curve (Supplemental Fig. 1).

### Iron content

Astrocyte media iron concentrations were measured by the ICP-AES method. Briefly, 1:1 volumes of media and 70% ultrapure nitric acid were added to glass tubes and incubated at 60 °C for 18 hours. Digested media was centrifuged at 10,000 g for 10 minutes at room temperature, and the supernatant fraction was collected and diluted with metal-free water. Iron concentration was determined ICP-AES against internal standards. Results were expressed as µg Fe/ml of media.

### ^55^Fe Uptake

Astrocytes were plated 20,000 cells/cm^2^ on Matrigel-coated plates. The following day, cells were co-incubated with either 50 nM Aβ or untreated control and 5 μCi of ^55^Fe-NTA. After 72 hours of incubation, the cells were washed x3 with PBS and then dissolved in 100 μl 0.2 M NaOH for 30 minutes. Once fully dissolved, the solution was collected and added to scintillation vials along with 10 mL CytoScint scintillation cocktail (MP Biosciences). Samples were counted using the Hidex 300 SL (LabLogic) for three minutes each. Blank tube values were subtracted from final counts to correct for background counts. Blank Matrigel-coated wells were simultaneously incubated with either 50 nM Aβ or untreated control and 5 μCi of ^55^Fe-NTA and further processed to subtract any Matrigel-captured ^55^Fe.

### CCK8 Mitochondrial Activity

Astrocytes were plated 20,000 cells/cm^2^ on Matrigel-coated plates. The following day, cells were co-incubated with either 50 nM Aβ or untreated control and 1:50 CCK8 reagent (Abcam, ab228554) and astroglial media. CCK8 uses a tetrazolium salt that is reduced in the presence of active mitochondria; thus it was used as a measure of metabolic activity. After 72 hours of incubation, media was collected and absorbance was read at 460 nm. To normalize the results to cell count, the cells were washed with PBS and lifted using TrypLE Express Enzyme (Thermo, 12604013). The cell solution was spun down and the pellet resuspended. Cell count was obtained using trypan blue and the Countess II Cell Counter (Thermo). There was no change to cell count with control or Aβ treatment.

### Immunoblotting

Samples were loaded onto a 4-20% Criterion TGX Precast Protein Gel (Bio-Rad)^11^. Protein was transferred onto a nitrocellulose membrane and probed for Fpn (Alpha Diagnostics, MTP11-S, 1:1000), DMT1 (Millipore, ABS983, 1:1000), Heph (Santa Cruz, SC-365365, 1:1000), TfR (Santa Cruz, sc-65882, 1:250), NEP-1 (ProteinTech, 18008-1-AP, 1:1000), APP (ProteinTech, 60342-1-Ig, 1:1000), IRP1 (Cell Signaling, 20272, 1:1000), IRP2 (Cell Signaling, 37135, 1:1000), FTH (Cell Signaling, 4393 1:1000), FTL (Abcam, ab69090 1:1000), Cytochrome C (ProteinTech, 66264-1-Ig, 1:1000), TOM20 (ProteinTech, 66777-1-Ig, 1:1000), and beta-actin (Sigma, a5441, 1:1000) or cyclophilin B (Abcam, ab16045, 1:1000) as a loading control. Corresponding secondary antibody conjugated to HRP was used (1:5000, GE Amersham) and bands were visualized using ECL reagents (Perkin-Elmer) on an Amersham Imager 600 (GE Amersham). Cellular lysate samples were normalized to cyclophilin B protein as a loading control, and then subsequently normalized to an untreated control sample within each experiment. Membrane protein samples were stained with Ponceau S and normalized to total protein as a loading control.

### Experimental Design and Statistical Analysis

Astrocytes were exposed to 50 nM Aβ or nothing (control) in media for 72 hours, after which the conditioned media (CM) was collected. ECs were culture in Transwell inserts. Aβ and control CM were placed in the basal chamber to measure radioactive iron transport from ECs. The presence of known iron stimulators was tested in the CM via ELISA and urea gel shift assay with control CM and/or media alone serving as controls. The effects of these molecules on iron transport from ECs was also tested as before with no molecule treatment serving as control. Mechanistic changes were investigated in astrocytes and ECs using western blot and functional assays (CCK8 and ^55^Fe uptake) with no treatment serving as controls.

Statistical analyses were performed using Prism 9.5 software (Graphpad Software Inc.). Data from at least three independent biological replicates were averaged and are expressed as the mean ± standard error of the mean (SEM). One-way ANOVA with Tukey post-hoc analysis, two-way ANOVA with Sidak’s post hoc analysis, or unpaired t-tests were used to evaluate for statistical significance where appropriate. A p-value <0.05 was considered significant.

## Results

### Media from Aβ-treated astrocytes increases iron transport from ECs

To first determine if Aβ-treated astrocytes could stimulate the transport of iron from ECs, astrocytes were treated with 50 nM Aβ for 72 hours to simulate the chronic nature of Aβ exposure. The control condition was no Aβ exposure. The conditioned media (CM) was collected from both control and Aβ-treated astrocytes. ECs were cultured onto Transwell inserts and control CM or Aβ CM was placed in the basal chamber to represent the brain-side. In the apical chamber, ^55^Fe-Tf and RITC-dextran were added, and at various intervals, aliquots were taken from the basal chamber to measure the transport of ^55^Fe-Tf and the leakage of RITC-dextran. ECs incubated with Aβ CM displayed a significant 3-fold increase of ^55^Fe-Tf transport over 24 hours (**p<0.01, Fig. 1A). Neither Control CM nor Aβ CM induced any barrier leakage (Fig. 1B).

**Figure 1:**
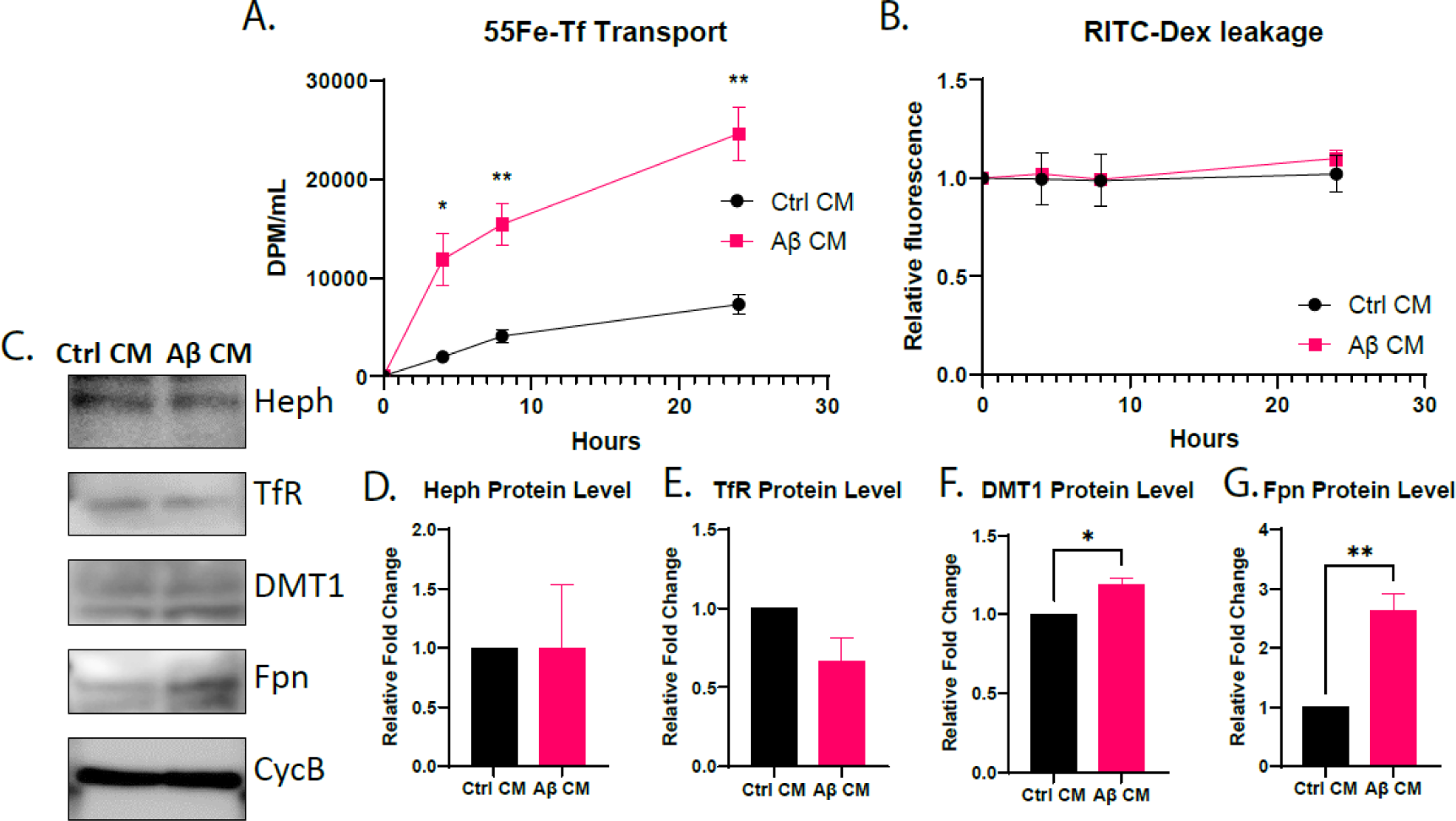
Modulation of iron transport from ECs by Aβ conditioned media from astrocytes. iPSC-derived astrocytes were treated with nothing (control) or 50 nM Aβ and the respective control (Ctrl) conditioned media (CM) and Aβ CM were collected after 72 hours. iPSC-derived ECs were cultured on bi-chamber plates and incubated with Ctrl CM or Aβ CM in the basal chamber and 10 μCi ^55^Fe-Tf and 1mg/ml RITC-dextran in the apical chamber. Aliquotes were taken from the basal chamber to measure ^55^Fe-Tf transport and RITC-dextran leakage. ECs incubated with Aβ CM had a 3-fold increase of ^55^Fe-Tf transport (**A**) with no change to monolayer permieability (**B**) compared to ECs exposed to Ctrl CM. ECs incubated with Ctrl CM and Aβ CM for 8 hours were collected for immunoblotting with all proteins normailized to cyclophilin B (CycB) (**C**). Hephaestin (Heph) (**D**) and transferrin receptor (TfR) (**E**) levels were unchanged in either condition. Divalent metal transporter (DMT1) (**F**) and ferroportin (Fpn) (**G**) levels were increased in ECs exposed to Aβ CM. n=3 for all experiments, means of biological replicates ± SEM were evaluated for statistical significance using two-way ANOVA with Sidak’s posttest for significance (A-B) or using unpaired t-test (D-G). *p<0.05, **p<0.01

To further investigate changes in iron regulatory proteins resulting in an increase of iron transport, ECs were collected for immunoblotting after basal incubation of control CM and Aβ CM for 8 hours. Ferroportin (Fpn), the only known iron exporter protein, was increased by nearly 3-fold in ECs incubated with Aβ CM (**p<0.01, Fig. 1C). Divalent metal transporter (DMT1), which transports from the endosome into the cytosol was also increased (*p<0.05, Fig 1C). Hephaestin (Heph) and transferrin receptor (TfR) were unchanged (Fig. 1C).

### Aβ-conditioned media contains known iron transport stimulators

In observing increased iron transport from ECs and corresponding changes to iron transport proteins after Aβ CM incubation, we examined multiple components of the control CM and Aβ CM previously demonstrated to have an impact on iron transport in ECs^9, 22–24^. First, the levels of Aβ_4222_ were measured in control CM, Aβ CM, and media alone (never exposed to cells) with 50 nM Aβ added to determine how much Aβ remained in the CM after incubation with astrocytes. Aβ CM contained about 10% less Aβ than the media where 50 nM Aβ was added (*p<0.05, Fig. 2A), which is consistent with literature suggesting that astrocytes participate in Aβ clearance^25^. Iron levels^9^ in the control CM, Aβ CM, and media alone were determined using ICP-MS. Aβ CM contained 50% less iron than control CM (**p<0.01) and was below media without cells (Fig. 2B), indicating an increase of iron uptake by the astrocytes. To accompany the decrease of media iron content, we hypothesized the percentage of apo-Tf (iron free) would be similarly increased. To determine the percentage of apo-Tf in the media samples, we used a urea gel shift assay (Fig. 2C). The band intensity of both apo- and holo-Tf were determined and the ratio was compared to the standard curve (supplemental Fig. 1) to determine the percentage of apo-Tf. Aβ CM contained twice as much apo-Tf compared to control CM (****p<0.0001, Fig. 2D). Additional suspected iron stimulators were measured by ELISA. Soluble amyloid precursor protein-α (sAPP-α)^23^ in control CM and Aβ CM were measured and found to be elevated 2-fold in Aβ CM (**p<0.01, Fig. 2E). As a measure of general cytokine production, IL-6^24^ was elevated 1.5-fold in Aβ CM compared to control CM (****p<0.0001, Fig. 2F).

**Figure 2:**
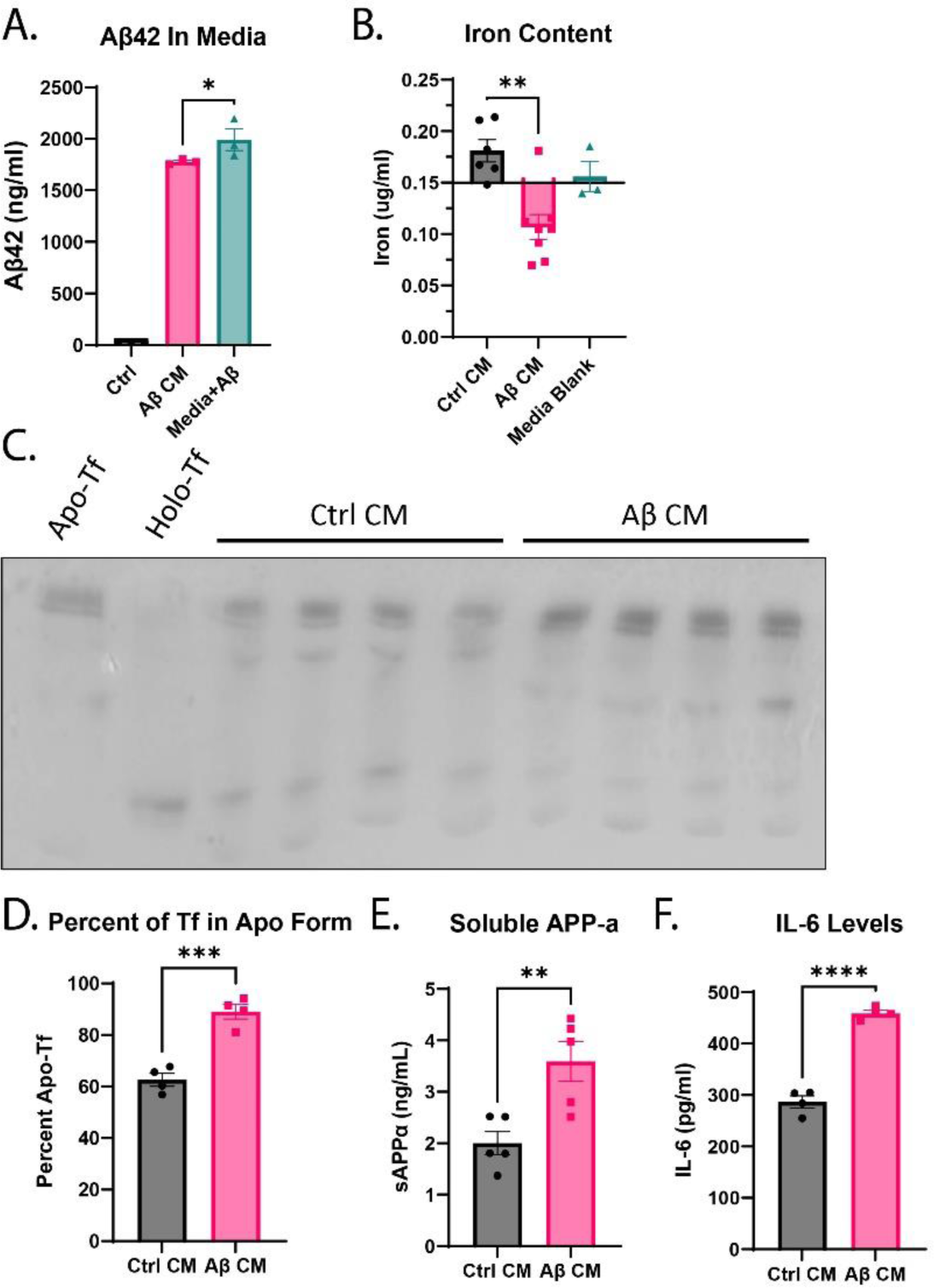
Potential iron transport stimulators found in conditioned media. iPSC-derived astrocytes were treated with control or 50 nM Aβ and the respective control (Ctrl) conditioned media (CM) and Aβ CM were collected after 72 hours for further analysis. The amount of Aβ_42_ still present in the media was assessed using ELISA (**A**). 50 nM Aβ was added to astrocyte media and incubated in an empty 6-well plate for 72 hours to compare. In the Aβ CM, about 10% of the Aβ has been degraded by the astrocytes, though a large amount remains present in the Aβ CM (**A**). Iron content of the media was determined using ICP-MS (**B**). Astrocyte media not exposed to cells was used as a blank measurement. Aβ CM contained about 50% less iron than Ctrl CM and less than the baseline media (**B**). The percentage of apo-Tf in the media was determined using a urea gel shift assay. The band intensity of both apo- and holo-Tf bands were measured and the ratio of apo:holo was calculated. Using the standard curve made with known ratios of apo:holo, the percentage of Tf in the apo form was calculated (**C**). Aβ CM contains about 30% more apo-Tf than Ctrl CM (**D**). Levels of soluble amyloid precusor protein-α (sAPP-α) were measured using ELISA (**E**). Aβ CM contained almost 2-fold more sAPP-α than Ctrl CM (**E**). Levels of IL-6 as a marker of general inflammatory response were measured using ELSIA (**F**). Aβ CM contained 50% more IL-6 than Ctrl CM (**D**). n=3 to 8 for all experiments, means of biological replicates ± SEM were evaluated for statistical significance using one-way ANOVA with Tukey’s posttest for significance (A-B) or using unpaired t-test (C, D, F). *p<0.05, **p<0.01, ****p<0.0001

### Astrocytes treated with Aβ display increased iron uptake and mitochondrial activity

After observing decreased iron content and increased percentage of apo-Tf in Aβ CM, we tested the hypothesis that the decrease in media iron content was due to increased iron uptake and mitochondrial activity^26^ after exposure to Aβ. Astrocytes plated in equal densities were incubated with ^55^Fe-NTA with or without 50 nM Aβ and collected after 72 hours to measure ^55^Fe uptake. Astrocytes treated with Aβ took up nearly double the amount of ^55^Fe than control (**p<0.01, Fig. 3A). In order to test mitochondrial activity, we employed a CCK8 assay. The CCK8 reagent contains a tetrazolium salt that is reduced by active mitochondria resulting in a colorimetric change. Astrocytes were incubated with CCK8 reagent with or without 50 nM Aβ, and after 72 hours, the media was analyzed and number of adherent cells was counted to normalize CCK8 absorbance. CCK8 absorbance increased by almost 2-fold after astrocytes were treated with Aβ (**p<0.01, Fig. 3B), indicating an increase of mitochondrial activity. Because the CCK8 assay is often used to measure cell viability we wanted to confirm no change in cell viability with or without Aβ treatment. We measured LDH levels in the control CM and Aβ CM, and there was no change in control CM or Aβ CM (Fig. 3C), indicating no increase of cell death. To confirm our functional findings, astrocytes incubated with 50 nM Aβ or control media for 72 hours were collected to assess various protein level changes via immunoblotting. Neprilysin (NEP-1) is an astrocytic Aβ-degrading enzyme and is upregulated in response to Aβ^27^ compared to control (*p<0.05, Fig. 3E). Additionally, APP, the precursor to sAPP-α, was decreased after Aβ treatment (*p<0.05, Fig. 3F). When intracellular iron levels increase, either Iron response protein (IRP) 1 or IRP2 is reduced in order to regulate iron export and storage, though rarely both^28^. In astrocytes after Aβ treatment, while not statistically significant, IRP1 decreased by 20% (Fig. 3G) and IRP2 was unchanged (Fig. 3H). Consistent with these findings, ferritin heavy chain (FTH) and ferritin light chain (FTL), both proteins that form ferritin to store intracellular iron, are increased by 2-fold in astrocytes treated with Aβ, though only FTL reaches statistical significance (*p<0.05, Fig. 3I-J). Cytochrome C is an iron complex protein used in the electron transport chain within mitochondria, and TOM20 is a receptor found on the outer mitochondrial membrane. Both cytochrome C and TOM20 are often used as mitochondrial markers, and both are increased in astrocytes after Aβ treatment (*p<0.05, Fig. 3K-L).

**Figure 3:**
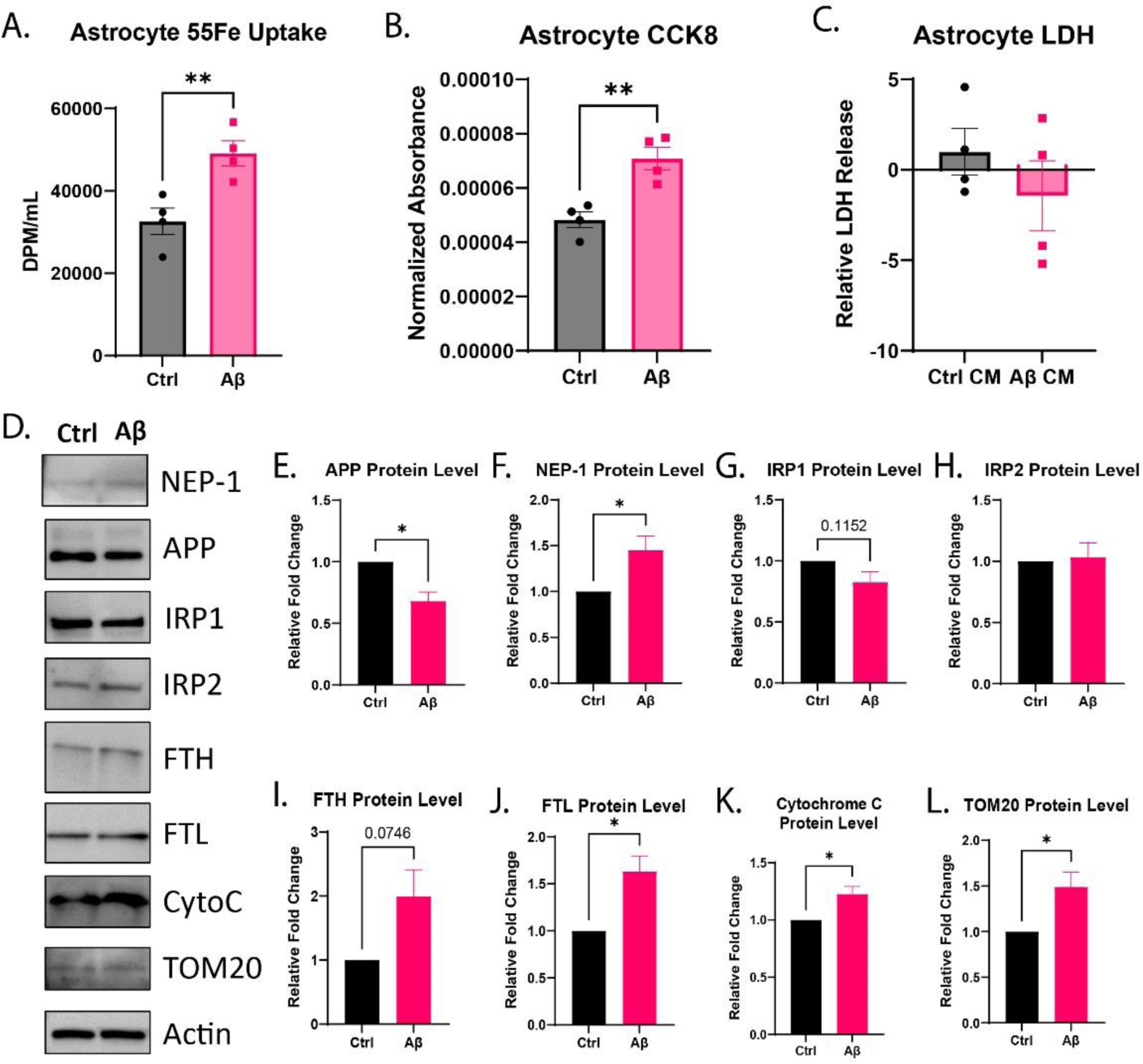
Aβ-induced changes in astrocytes. iPSC-derived astrocytes were treated with nothing (control, Ctrl) or 50 nM Aβ for 72 hours. To measure iron uptake, cells were co-incubated with 5 μCi of ^55^Fe-NTA and Aβ, and after 72 hours, the cells were dissolved and collected for liquid scintilation counting (**A**). Astrocytes treated with Aβ had a significant increase of ^55^Fe uptake (**A**). To measure mitochondrial activity, cells were co-incubated with CCK8 reagent and Aβ, and after 72 hours, the media was collected to measure CCK8 reduction and the cells were collected for cell count (**B**). Astrocytes treat with Aβ had a signiciant increase of mitochondrial activity normailized to cell count (**B**). To measure cell viability after Aβ treatment, LDH was measured in the control (Ctrl) CM and Aβ CM and found to have no change (**C**). To determine molecular changes to confirm the functional findings, cells were collected for immunoblotting after Aβ treatment (**D**). Changes to IRP1, FTH, and FTL levels supported increased cellular iron content (**E, I, J**). Increases of cytochrome C and TOM20 levels supported increased mitochondrial activity with Aβ treatment (**K-L**). APP levels were decreased supporting the increase of sAPP-α in the Aβ CM (**G**). NEP-1 levels were increased with Aβ treatment, supporting astrocytic response to Aβ (**H**). n=3 to 4 for all experiments, means of biological replicates ± SEM were evaluated for statistical significance using unpaired t-test. *p<0.05, **p<0.01

### sAPP-α alone does not increase iron transport from ECs

Our group has previously demonstrated that iron deficient conditions and apo-Tf stimulate iron transport from ECs^8, 9, 11^, and sAPP-α has been reported to increase iron release from ECs^23^. To determine if either apo-Tf or sAPP-α could mimic the iron transport stimulation by Aβ CM, ECs were cultured onto Transwell inserts and 0.25 μM apo-Tf (physiological in CSF^29^), 0.1 nM sAPP-α (concentration in Aβ CM), apo-Tf and sAPP-α, or control was placed in the basal chamber. Again, the apical chamber contained ^55^Fe-Tf and RITC-dextran, and at hourly intervals, aliquots were taken from the basal chamber to measure the transport of ^55^Fe-Tf and assess the integrity of the tight junctions with RITC-dextran. As previously shown, apo-Tf stimulated ^55^Fe-Tf transport compared to control (**p<0.01, Fig. 4A), however, sAPP-α failed to stimulate iron transport alone (Fig. 4A). The combination of sAPP-α and apo-Tf did not further increase ^55^Fe-Tf transport than apo-Tf alone (*p<0.05, Fig. 4A), indicating that apo-Tf was driving the response. Neither apo-Tf nor sAPP-α increased RITC-dextran leakage (Fig. 4B). Fpn levels were examined to determine if either apo-Tf or sAPP-α altered the levels of Fpn in ECs similar to Aβ CM did, and Fpn levels remained consistent across experimental conditions (Fig. 4C-D).

**Figure 4:**
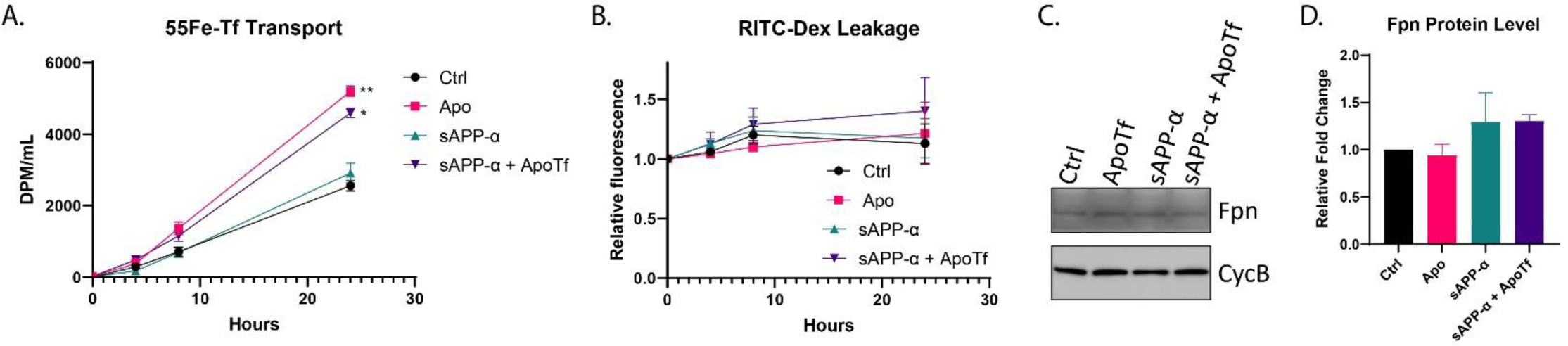
Lack of effect by soluble APP-α on iron transport from ECs. iPSC-derived ECs were cultured on bi-chamber plates and incubated with 0.25 μM apo-Tf and/or 0.1 nM sAPP-α in the basal chamber and 10 μCi ^55^Fe-Tf and 1mg/ml RITC-dextran in the apical chamber. Aliquotes were taken from the basal chamber to measure ^55^Fe-Tf transport and RITC-dextran leakage. Apo-Tf and the combination of apo-Tf and sAPP-α increased ^55^Fe-Tf transport but sAPP-α alone did not (**A**). All conditions did not impact monolayer permieability (**B**). ECs incubated with apo-Tf and/or sAPP-α for 8 hours were collected for immunoblotting for Fpn levels, which were normailized to cyclophilin B (CycB) (**C**). No experimental condition increased ferroportin (Fpn) (**D**) levels. n=3 for all experiments, means of biological replicates ± SEM were evaluated for statistical significance using two-way ANOVA with Sidak’s posttest for significance (A-B) or using one-way ANOVA with Tukey’s posttest for significance (D). *p<0.05, **p<0.01

### Traditional inflammatory cytokines do not elicit changes in iron transport from ECs

Numerous studies have shown cytokine-induced EC monolayer integrity dysfunction^24, 30–32^ and that cytokines or neuroinflammation can impact iron uptake in the brain^33, 34^. To determine if classically AD-elevated cytokines^35^ could mimic the iron transport stimulation of Aβ CM, ECs were cultured on Transwell inserts and combinations of 50 nM Aβ, 100 ng/mL IL-6, 50 ng/ml tumor necrosis factor α (TNF-α), and/or 100 ng/ml interleukin-1β (IL-1β), or control were placed in the basal chamber, and ^55^Fe-Tf and RITC-dextran were placed in the apical chamber. Aliquots were taken from the basal chamber to measure the transport of ^55^Fe-Tf and the leakage of RITC-dextran. Neither solo exposures of Aβ, IL-6, TNF-α, or IL-1β nor combinations increased ^55^Fe-Tf transport (Fig. 5A and C) or RITC-Dextran leakage (Fig. 5B and D). Fpn levels were examined to determine if any cytokine treatment or Aβ increased EC Fpn, and Fpn levels remained consistent across experimental conditions (Fig. 5E-F).

**Figure 5:**
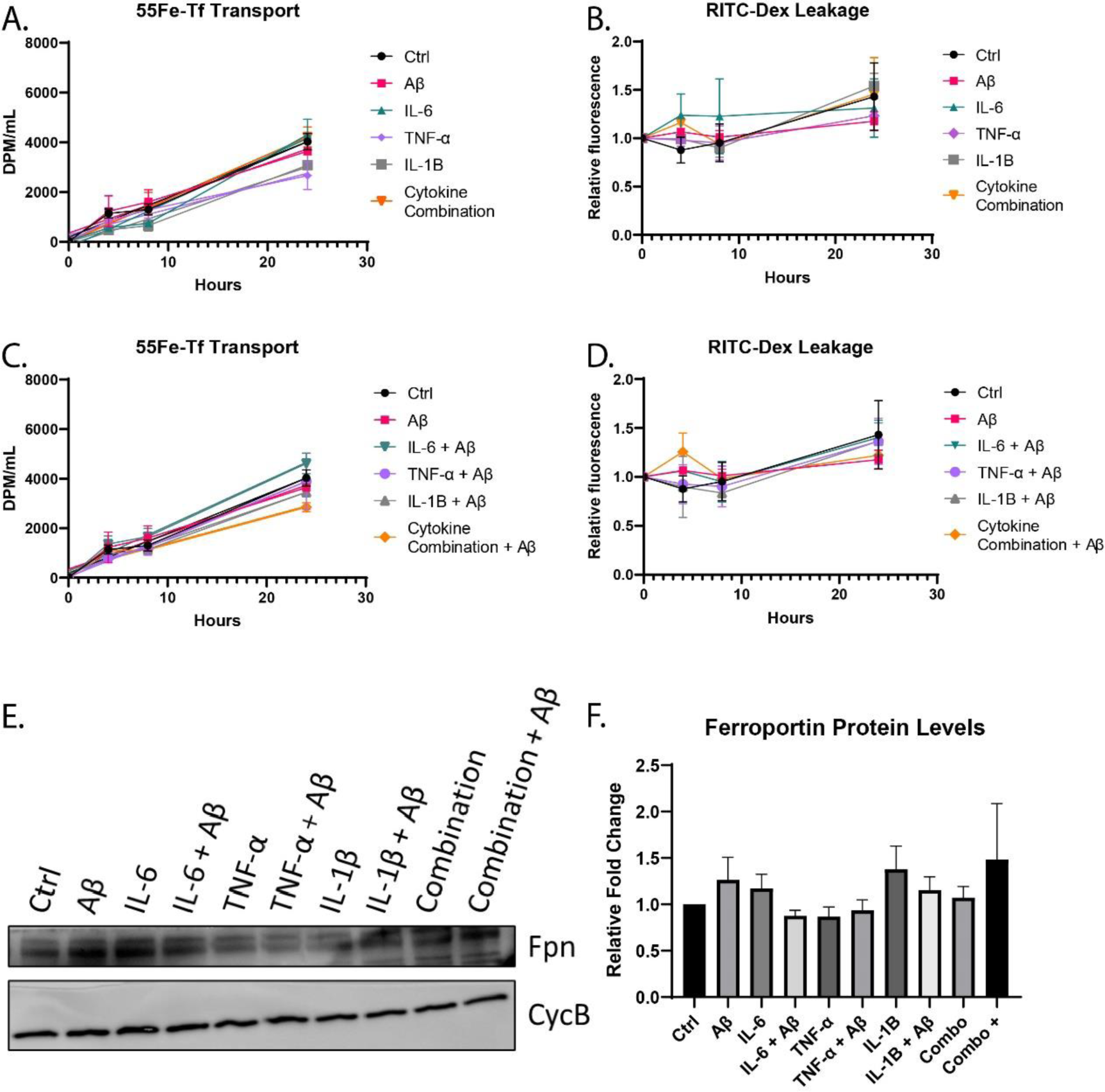
Lack of effect by cytokines on iron transport from ECs. iPSC-derived ECs were cultured on bi-chamber plates and incubated with combinations of 50 nM Aβ, 100 ng/mL IL-6, 50 ng/ml TNF-α, and/or 100 ng/ml IL-1β in the basal chamber and 10 μCi ^55^Fe-Tf and 1mg/ml RITC-dextran in the apical chamber. Aliquotes were taken from the basal chamber to measure ^55^Fe-Tf transport and RITC-dextran leakage. None of the cytokines nornor Aβalone or in combination impacted ^55^Fe-Tf transport (**A and C**) or to monolayer permieability (**B and D**). ECs incubated with Aβ and cytokines for 8 hours were collected for immunoblotting to for Fpn levels normailized to cyclophilin B (CycB) (**E**). Noexperimental condition increased ferroportin (Fpn) (**F**) levels. n=3 for all experiments, means of biological replicates ± SEM were evaluated for statistical significance using two-way ANOVA with Sidak’s posttest for significance (A-D) or using one-way ANOVA with Tukey’s posttest for significance (F).

## Discussion

This study suggests a possible mechanism for excessive iron accumulation in response to early AD Aβ deposition. Herein, we demonstrate that astrocytes treated with low levels of Aβ, consistent with early AD pathology, stimulate the transport of iron from ECs at the BBB (Fig. 6). An investigation into the media components for known stimulators of iron release found that Aβ CM from astrocytes contained less iron and a higher percentage of apo-Tf than control CM. Additionally, levels of sAPP-α and inflammatory cytokines such as IL-6 were elevated in Aβ CM. In response to Aβ exposure, astrocytes increased their mitochondrial activity which was consistent with increased iron uptake. The increase iron need by the astrocytes is accompanied with increased release of apo-Tf. When ECs were incubated with apo-Tf, iron transport was increased, as previously demonstrated^8, 9, 11^. Other reported iron release stimulators, such as sAPP-α, Aβ, and inflammatory cytokines, did not mimic the effects of Aβ CM on iron transport in ECs. These findings suggest the mechanism for the initiation of excessive brain iron accumulation in AD starts with Aβ creating an iron deficient environment that promotes iron uptake mechanisms, leading to misappropriation of the astrocyte-EC iron regulatory signally and subsequent iron accumulation.

**Figure 6:**
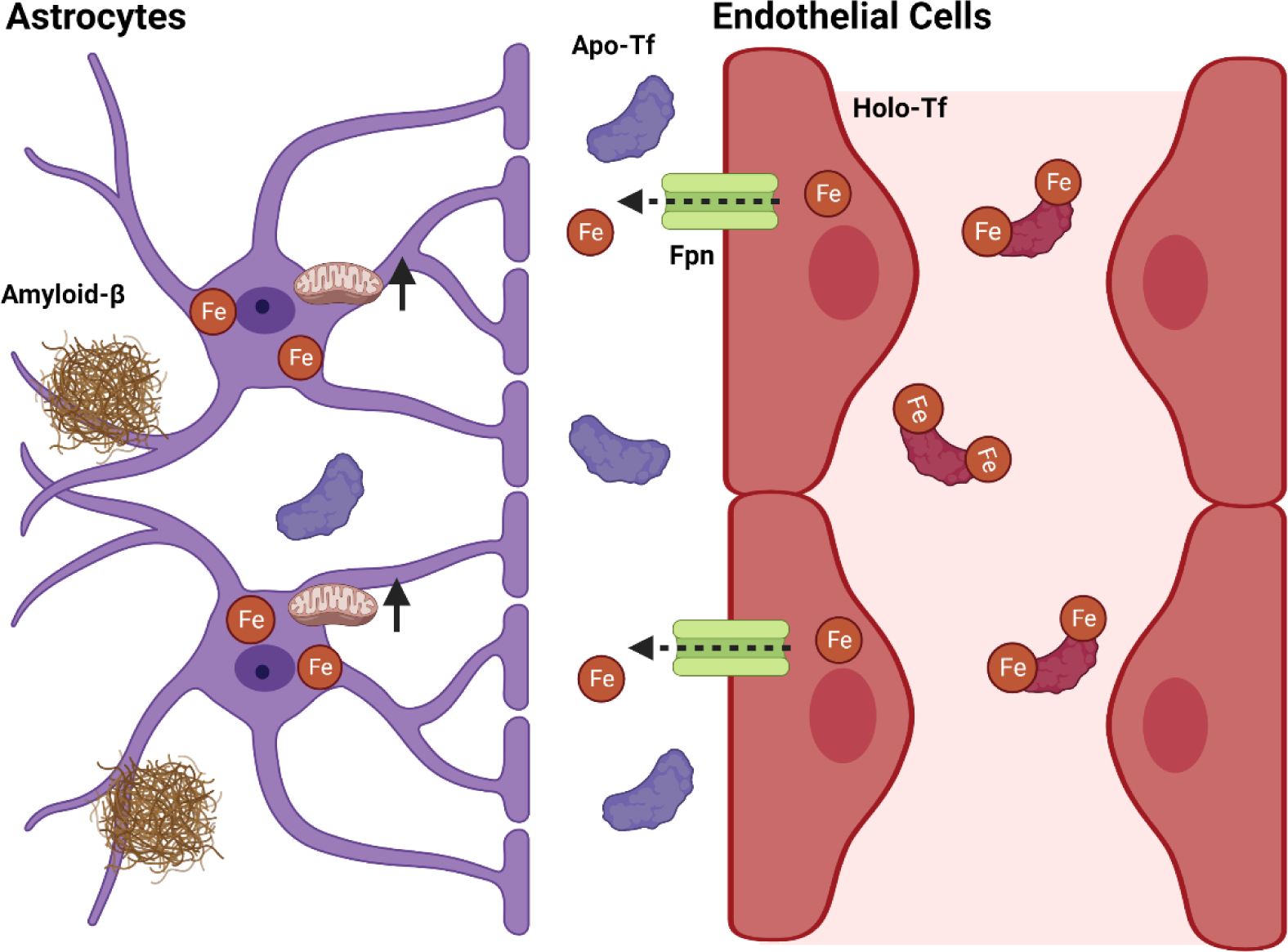
Summary Model. In response to amyloid-β (Aβ), astrocytes increase their mitochondrial activity, partially in response to an increase of energetic needs due to Aβ clearance. This is accompanied by an increase of iron uptake and consumption, leaving an iron deficient enviornment in the extracellular space and elevated levels of apo (iron free)-transferrin (Tf). Apo-Tf stimulates iron release from endothelial cells (ECs), resulting in an increase of iron transport across the blood-brain barrier.

The process of iron transport at ECs is highly regulated, which allows for proper neural functioning^36^. In order to cross the BBB, holo-Tf binds to TfR on the luminal membrane (blood side). The complex is endocytosed and the iron is reduced and transported out of the endosome by DMT1. Once free in the cell, iron is either used in the labile iron pool, stored in ferritin or exported via Fpn. Fpn is the major focus for most studies on iron release regulators. Heph stabilizes Fpn and aids in iron export by oxidizing the free iron released into the extracellular fluid and enables binding to apo-Tf^37^. In the present study, ECs exposed to Aβ CM significantly increased iron transport with no change to EC monolayer permeability. These data suggest that the ECs themselves were not damaged or leaky, but rather the process of iron transport was engaged. When proteins involved in iron transport were examined, DMT1 and Fpn were found to be increased in ECs exposed to Aβ CM, which is consistent with an increase of iron transport and release. TfR was unchanged, but it is noteworthy that TfR is found on both the luminal and basolateral membranes^9^ and our technique does not differentiate between the two TfR populations. Taken together, these data suggest that a component in Aβ CM signals to ECs to increase iron transport.

Our group has studied at length how apo-and holo-Tf act as signals of brain iron status in order to modulate iron release from ECs with corresponding molecular changes^8, 9, 13, 37^. Apo-Tf signals an iron deficient environment and stimulates iron release from ECs, whereas holo-Tf signals an iron saturated environment and suppresses iron release. In line with reduced iron release, we have shown that ECs incubated with holo-Tf have reduced Fpn levels^9, 37^. Recently, LeVine *et al.* found that iron deficiency, iron transport, and mitochondrial related processes were all upregulated in AD patient tissue^7^. Here, we found that Aβ CM contained significantly less iron than control CM. In line with this finding, we also discovered that of the Tf present in Aβ CM, the larger percentage of it was apo-Tf compared to control CM. Furthermore, when ECs were incubated with apo-Tf in the basal chamber, iron transport was increased, supporting our previous demonstration^8, 9^ and suggesting that an increase of apo-Tf in Aβ CM is a substantial contributor to the observed increase in iron transport.

One possible explanation for a decrease of iron in Aβ CM from astrocytes would be increased iron uptake due to metabolic activity. Astrocytes take up large amounts of iron and can increase their iron content by 9-fold before viability is affected^38^. Once iron is taken up by astrocytes, the iron is either stored in ferritin or incorporated into iron-sulfur clusters, which are crucial electron transfer molecules in mitochondrial respiration such as cytochrome C^39^. Studies have shown higher amounts of Aβ can negatively impact astrocytic metabolism^25, 26^. Here, we found that astrocytes treated with 50 nM Aβ (widely considered physiological^40^) display increased iron uptake and mitochondrial activity compared to control. Additionally, there was no change in cell viability with or without Aβ treatment, suggesting the increased iron content is not yet detrimental to the cells. An examination of molecular changes in the astrocytes corroborated our functional findings. Proteins associated with increased iron storage (FTH and FTL) and mitochondria density (TOM20 and cytochrome C) were increased in astrocytes treated with 50 nM Aβ. While there are multiple possibilities for increased energetic needs in response to Aβ, two measured here were APP and NEP-1. Decreased APP levels suggest an increase of APP cleavage, and increased NEP-1 levels suggest an increase of Aβ clearance. Taken together, these data suggest that, in response to low levels of Aβ, astrocytes increase their mitochondrial activity, leading to increased iron consumption and an iron deficient extracellular environment. All of these processes are functions of normal debris clearance and iron transport regulatory mechanisms; however, the chronic nature of AD likely leads to chronic iron build-up and damage over time.

sAPP-α is another potential iron release stimulator secreted by astrocytes. APP is a transmembrane protein most commonly known for its role in Aβ formation, though also plays an important role in iron release from ECs^41, 42^. Amyloidogenic processing occurs when Aβ-secretase incorrectly cleaves APP, thus producing insoluble pathogenic Aβ^43^. Non-amyloidogenic processing occurs when α-secretase cleaves APP and produces sAPP-α^43^. Numerous groups have shown that APP is required for Fpn function and stability in the plasma membrane^41^ through direct binding to Fpn^42^. The Fpn targeting peptide sequence present within APP is also present in sAPP-α^23^. McCarthy *et al.* demonstrated that sAPP-α similarly binds to Fpn and 10 nM treatment of sAPP-α induces iron release from ECs, though not due to ferroxidase activity like Heph^23^. In our present study, we found that treatment with 50 nM Aβ increased levels of sAPP-α secreted from astrocytes. This observation combined with reduced levels of membrane APP in astrocytes treated with Aβ suggests that Aβ stimulates non-amyloidogenic processing of APP leading to increased sAPP-α production. Our study did not find an increase of iron transport from ECs associated with sAPP-α alone, but the Aβ CM only contained 0.1 nM of sAPP-α compared to the 10 nM required to induce iron efflux^23^, suggesting that sAPP-α is not a cause for the Aβ CM-induced increase in iron transport from ECs seen in this study.

Another potential inducer of iron release at the BBB following activation of astrocytes is the release of inflammatory cytokines which has been shown in response to Aβ exposure with both *in vitro*^44^ and *in vivo* models^45^. In AD patient CSF, increases of inflammatory cytokines, IL-6 and TNF-α, positively correlate with AD pathology^46^ and with worsening cognitive impairment^47, 48^. Numerous studies have shown dysfunction in BBB permeability after exposure to inflammatory cytokines^24, 30–32^. Notably, de Vries *et al.* found that IL-6, TNF-α, and IL-1β reduced EC monolayer integrity by about 60% in rat cerebral ECs^24^. Other studies have shown increases of Fpn and TfR, which are indicative of increase iron transport, in response to Aβ *in vivo*^49^. Contrary to these previous studies, Kim *et al* showed that, in order to negatively impact EC monolayer permeability, reactive astrocytes needed direct contact with the ECs and that single doses of various inflammatory cytokines did not impact ECs alone^18^. In our present study, IL-6, TNF-α, and IL-1β in single doses and combination doses with 50 nM Aβ do not have any impact on iron transport or monolayer permeability compared to control. In addition, none of these experimental conditions influenced Fpn levels in ECs. Our findings support those of Kim *et al.* and indicate that IL-6, TNF-α, IL-1β, or Aβ cannot account for the increased iron transport from ECs after incubation with Aβ CM.

The narrative of iron in AD has long focused on how iron contributes to mid-stage disease pathology and iron overload. Our data uniquely point to an early-stage disease initiation of excessive iron accumulation. This study is the first to start to decipher the process of how the normal communication between astrocytes and ECs can be misappropriated by a toxin to initiate dysregulation of brain iron acquisition. We show that in response to Aβ, astrocytes increase their mitochondrial activity and iron consumption, resulting in the secretion of iron release signals (apo-Tf) to ECs. These data build on our demonstration of the role of apo- and holo-Tf in regulating brain iron uptake and expand it into a pathological setting. Understanding how this normal response to a pathological substance result in excessive brain iron accumulation in AD may provide a possible therapeutic intervention point to mitigate the downstream damage of unchecked brain iron accumulation.

## Acknowledgements

The authors would like to thank Dr. Irina Elcheva and Stem Cell and Regenerative Program for their support. Research reported in this publication was supported by the National Center for Advancing Translational Sciences of the National Institutes of Health Award Number TL1TR002016. The content is solely the responsibility of the authors and does not necessarily represent the official views of the NIH. Funding for this study was provided by NIH R01NS113912-04 (JRC) and NIH R21AG064486-02 (JRC) and SLB was supported by NIH TL1TR002016 (SLB).

## Abbreviations

- Alzheimer’s disease (AD)
- Amyloid-β (Aβ)
- Amyloid precursor protein (APP)
- Blood-brain barrier (BBB)
- Conditioned medium (CM)
- Divalent metal transporter (DMT1)
- Endothelial cells (ECs)
- Ferritin heavy chain (FTH)
- Ferritin light chain (FTL)
- Ferroportin (Fpn)
- Hephaestin (Heph)
- Interleukin-1β (IL-1β)
- Interleukin-6 (IL-6)
- Iron response protein (IRP)
- Lactate dehydrogenase (LDH)
- Neprilysin (NEP-1)
- Soluble APP-α (sAPP-α)
- Transferrin (Tf)
- Transferrin receptor (TfR)
- Tumor necrosis factor α (TNF-α)

---

**Supplemental Figure 1:**
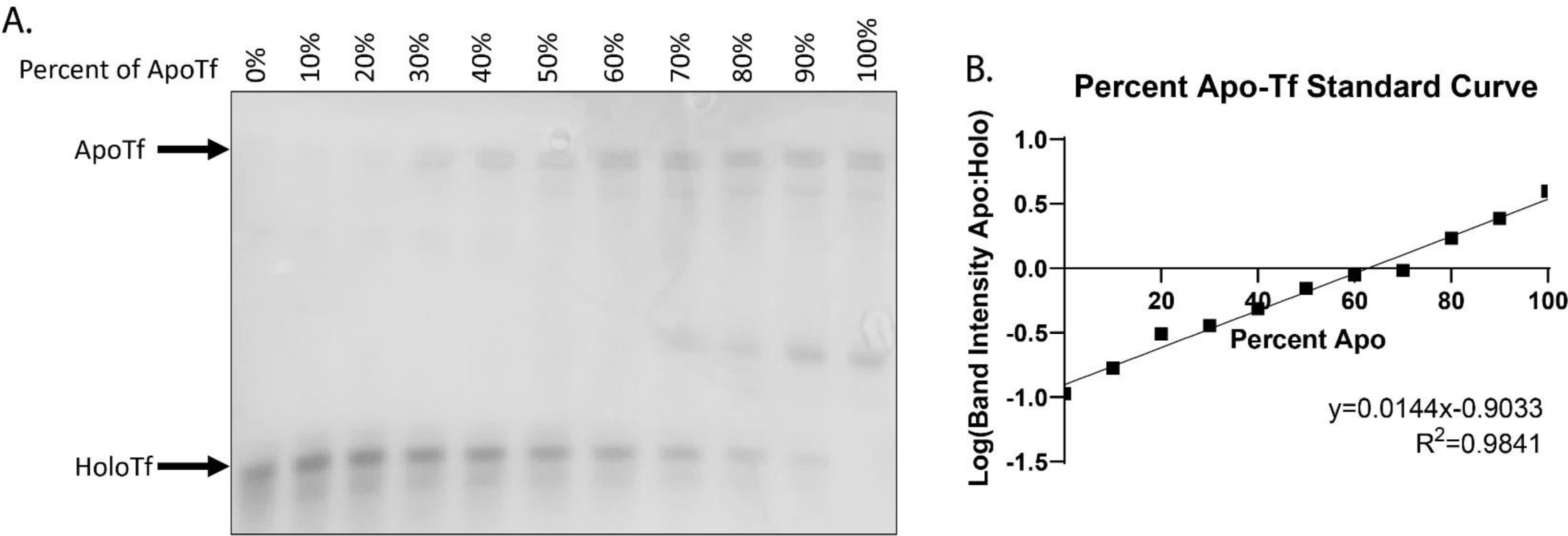
Urea Gel Shift Tf Standard Curve. In order to more accurately determine the percentage of apo-Tf present in experimental CM samples, a standard curve was created (**A**). Set percentages of apo- and holo-Tf were mixed together. Five μg of protein was run on TBE-urea gels and Tf bands were visualized using total protein stain. Because urea binds to the available iron binding sites on apo-Tf, apo-Tf runs slower through the gel compared to holo-Tf. This results in apo- and holo-Tf reliably separating as shown here (**A**). The band intensity of both apo- and holo-Tf were determined and the ratio of apo:holo was calculated. After performing the experiment three times, the ratios were averaged together. The log of the band intensity ratio was plotted against the known percent of apo-Tf in the solution to create the standard curve (**B**). The equation of the linear regression was used to determine the unknown percent of apo-Tf in CM samples based on the experimentally determined band intensity ratio.

